# Fecundity-immunity trade-off curvature modulates host evolutionary dynamics, pathogen propagation, and host abundance

**DOI:** 10.1101/2025.02.06.636764

**Authors:** Edgard B. Djahoui, Nicolas Loeuille, Rudolf P. Rohr

## Abstract

In most species, life history trade-offs generate conflicts between biological functions because of resource allocation constraints. During pathogenic infections, hosts can deploy various immune defenses, but these responses usually carry fitness costs, most often expressed as reduced fertility. This fecundity–immunity trade off has been widely studied for its evolutionary implications, yet its effects on other emergent properties —such as host abundance and pathogen spread— remain poorly understood. Here, we develop an SIR model incorporating three immune strategies —reduced host susceptibility, increased host recovery, and reduced pathogen virulence— and show that the evolutionary impact of the fecundity–immunity trade off depends on both its shape and the specific immune mechanism under selection. For reduced susceptibility and increased recovery, disease spread and host abundance evolve in opposite directions along evolutionary trajectories, such that when disease spread increases, host abundance decreases, and *vice versa*. For reduced virulence, however, this relationship is more complex and does not follow a simple pattern. Across all three mechanisms, trade offs with low cost intensity, whether concave or convex, favor the joint evolution of high fecundity and high immunity. In contrast, concave or convex trade offs incurring high cost consistently select for high fecundity at the expense of immunity. Trade offs with moderate cost intensity generate two alternative outcomes: one strategy with high fecundity and low immunity, and another combining low fecundity with enhanced immunity. Together, these findings underscore the importance of accounting for the specific immune mechanism evolving and the associated cost intensity when predicting the eco-evolutionary consequences of fecundity–immunity trade-offs.

**Highlights:**

- Impacts of host evolution on *R*_0_ and total host abundance *N* are rarely investigated
- Host evolution accessed under various components of a fecundity-immunity trade-off
- Weak fecundity-susceptibility / recovery trade-offs decrease *R*_0_ and increase *N*
- Strong fecundity-susceptibility / recovery trade-offs increase *R*_0_ and decrease *N*
- More complex patterns are observed for the fecundity-virulence trade-off

## 1 Introduction

In host–pathogen interactions, evolutionary changes usually take the form of co-evolution, in which adaptations in one partner generate counter-adaptations in the other. However, evolution may also be dominant in only one of the two species, affecting primarily either the host or the pathogen. These evolutionary dynamics often generate or modify system-level characteristics commonly referred to as emergent properties. Emergent properties are properties that appear or are modified in an interacting system and whose behaviors cannot be reproduced or predicted by considering any subset of the whole system (Johnson, 2006). One of the most important emergent properties in host-pathogen systems is pathogen virulence (ie, enhanced mortality rates during infection), which arises from the interplay of host and pathogen characteristics (Casadevall et al., 2011; Garcia-Solache et al., 2013). However, host–pathogen systems exhibit many other emergent properties, and investigating how evolutionary changes in the system affect them may provide deeper insight into the range of possible outcomes of such systems.

In host-pathogen systems, while host-only evolution under ecological constraints is possible, it can also arise from evolutionary pressures, especially from those imposed by the non-evolving pathogen. Indeed, following a pathogenic infection, the host usually mounts an immune response that activates a variety of defense mechanisms against the invading pathogen. However, activating the immune system is an energy-demanding process and may therefore interfere with other physiological functions. (Lochmiller and Deerenberg, 2000). Previous studies suggest that such costs are often expressed as reduced host fecundity or reproduction, leading to fecundity-immunity or reproduction-resistance trade-offs. These evolutionary constraints have been documented across a wide range of taxonomic groups, including insects, vertebrates, and plants (Züst and Agrawal, 2017; Buchanan et al., 2018; Rodrigues et al., 2021; Fedorka et al., 2004; Barbosa et al., 2016; Hurd et al., 1995; Ahmed and Hurd, 2006; Vézilier et al., 2012; Brokordt et al., 2019). For instance, in *Anolis carolinensis* lizards, food restriction drastically suppressed reproduction and immune function in the species, emphasizing shared resource allocation between these two metabolic processes (Husak et al., 2016). Since these host defense mechanisms or immune responses arise as a consequence of pathogen’s attack, they can be understood as emergent properties of host–pathogen interactions. Given their central importance, these host-side emergent properties have been extensively explored, assuming that only the host is evolving. Most often, this has been done through the lens of the ubiquitous fecundity–immunity trade-off, which in this context acts as the major evolutionary constraint driving host evolution.

Evolutionary dynamics of hosts *per se* have been largely investigated (reviewed in (Boots et al., 2008). These studies showed that fecundity-immunity trade-offs can generate and maintain resistant and susceptible strains in host polymorphism (Bowers et al. (1994); Antonovics and Thrall (1994); Boots and Haraguchi (1999); Boots and Bowers (1999, 2004); Miller et al. (2005)). (Best et al., 2015) later showed that the trade-off shape itself greatly affects this polymorphism. These previous studies also explored the partitioning of immunity through different defense mechanisms. (Restif and Koella, 2004), for instance, investigated the concurrent evolution of resistance and tolerance under different demographic and epidemiological conditions while (Best et al., 2008) investigated the emergence and maintenance of different tolerance mechanisms (mortality tolerance and sterility tolerance). These studies show that resistance and tolerance respond differently to changes in epidemiological and trade-off parameters, and that mortality tolerance increases parasite fitness, whereas sterility tolerance is neutral and can promote host polymorphism. (Best et al., 2017) further explored the evolution and coevolution of multiple defense strategies —avoidance, clearance, and tolerance— while (Boots and Best, 2018) analyzed the evolution of constitutive and induced defenses in relation to host mortality, crowding, and parasite virulence. Their findings indicate that high sterility selects for avoidance and clearance, with the potential coexistence of both traits, whereas low sterility favors tolerance. In addition, high virulence promotes stronger induced defenses despite the costs of immunopathology, ultimately resulting in elevated disease-induced mortality. Other studies examined the impact of host life span and other demographic and epidemiological characteristics on the optimum partitioning of innate vs acquired immunity (Boots et al., 2013; Donnelly et al., 2015), or on prevalence and proportion of immune (Miller et al., 2007), all using the fecundity-immunity trade-off as background constraint. Their results show that a higher abundance of infected individuals favors greater investment in innate resistance. Moreover, in the absence of acquired immunity, longer-lived populations invest more heavily in resistance, whereas in the presence of acquired immunity, the optimal level of resistance becomes dependent on host life span.

While these previous studies highlight various sides of evolutionary dynamics, they most often leave out their impacts at larger organization scales (population characteristics, epidemiological states). The present work aims at investigating such emergent properties, including disease spread, host productivity or abundance, and how such properties are related to one another along evolutionary trajectories. For instance, the level of pathogen virulence can influence the intensity of host defense, and *vice versa*, showing that some emergent properties can reciprocally affect one another. In van Boven and Weissing (2004), the effects of life span on investment in immunity, disease prevalence, and propagation were examined. The study, again, treated the fecundity-immunity trade-off as a background constraint and did not allow it to directly influence the emergent properties under investigation, most notably host total abundance, which was excluded from the analyses. Following their footsteps, we rely on an explicit and direct formulation of the trade-off, allowing host allocation strategies in fecundity and immunity to freely evolve under various trade-off shapes, and investigate how this affects host evolutionary strategies (immune VS fecund strategies), and how the resulting outcomes influence our emergent properties of interest.

Such a deep understanding of emergent properties is crucial for effective disease control. In epidemiology, for example, a key emergent property commonly used as a proxy for the intensity of pathogen propagation is the pathogen’s basic reproduction number or ratio, denoted by *R*_0_. *R*_0_ is defined as the number of secondary infections generated by a single infected individual in a completely susceptible host population(Martcheva, 2015; Lion and Metz, 2018). A pathogen can invade and generate an epidemic in its host population if and only if *R*_0_ *>* 1, with higher values of *R*_0_ implying larger outbreaks. Therefore, considerable effort has been devoted to identifying the environmental factors (biotic and abiotic) affecting *R*_0_ within given ecological and evolutionary contexts (Elderd and Reilly, 2014; Lion and Metz, 2018). However, most evolutionary studies on *R*_0_ have focused exclusively on pathogen evolution alone —typically through a virulence–transmission trade-off—, resulting in limited knowledge of how host evolution itself affects this emergent property of the system. It therefore becomes particularly interesting to assess how an explicit and direct fecundity–immunity trade-off, acting on host evolution, would influence pathogen propagation, as measured by *R*_0_. For example, in a host with strong immunity, susceptibility to infection may decrease as a direct effect. However, because immunity is subject to a resource allocation constraint, greater immune investment also reduces host reproduction, thereby decreasing the influx of susceptible individuals (assuming newborns are susceptible). These direct and indirect effects act synergistically to decrease *R*_0_.

Another emergent property of great interest is the total abundance of the host population. This property is highly sensitive to the interaction between pathogen attack and host defense mechanisms. It is well established how virulent pathogens can lead to host population decrease or collapse (Daszak et al., 2003; Robinson et al., 2010; Lebrun et al., 2022; Leendertz et al., 2006). Selection of efficient immune response, in principle, limits pathogen-associated damages. If such immune responses come at the cost of host fecundity, the link between the evolutionary dynamics and host abundance becomes non-trivial, as it depends on the balance between benefits (lower mortality) and costs (lower fecundity) of the evolving trait, hence on the trade-off shape. Such an understanding may be of central importance for the management of exploited species (eg, agricultural epidemics), a context in which total host abundance is directly linked to the final yield.

Our analysis will focus on studying the effects of fecundity-immunity trade-offs and their shapes (curvatures) on host evolutionary dynamics. We will study how such an evolution affects two emergent properties: (i) the basic reproduction number *R*_0_ as a proxy of disease propagation and eventual epidemiological state; (ii) the total host population density as a proxy of disease effects. This may also provide insights into the conditions that promote species coexistence or, alternatively, lead to the exclusion of the pathogen from the system. Then, we will examine the possible relationships that connect these two emergent properties under evolutionary changes. To achieve this, we use an epidemic model and investigate host evolution based on the adaptive dynamics framework. We decompose immunity into three components: host susceptibility to infection, host recovery, and the host’s ability to mitigate pathogen virulence. We then separately link each immunity component to fecundity, generating three fecundity-immunity trade-off scenarios, each exemplified by a specific biological system. These systems are subsequently analyzed under various trade-off shapes. For specific trade-off shapes where there is no or only very little cost associated with investing in either biological function, we expect the host to evolve toward higher levels of both fecundity and immunity simultaneously. Such an evolution should lead to an abundant host population and to low pathogen propagation. For trade-off shapes where investing in one biological function incurs a moderate cost in the other, we would then expect intermediate host abundance and pathogen propagation in the population. Trade-off shapes in which investing in one biological function strongly compromises the other are expected to exhibit two equally likely outcomes. The system could evolve either toward high fecundity and bad immunity, or toward low fecundity and good immunity, the intermediate positions becoming bad compromises between fecundity and immunity. High fecundity and poor immunity may then lead to high host abundance and high pathogen propagation, whereas low fecundity and good immunity could induce low host abundance and low pathogen propagation.

## 2 Material and methods

### 2.1 SIR model

We use a susceptible—infected—recovered (SIR) single host epidemic model depicted on figure 1 and given by the following set of differential equations:

**Figure 1.**
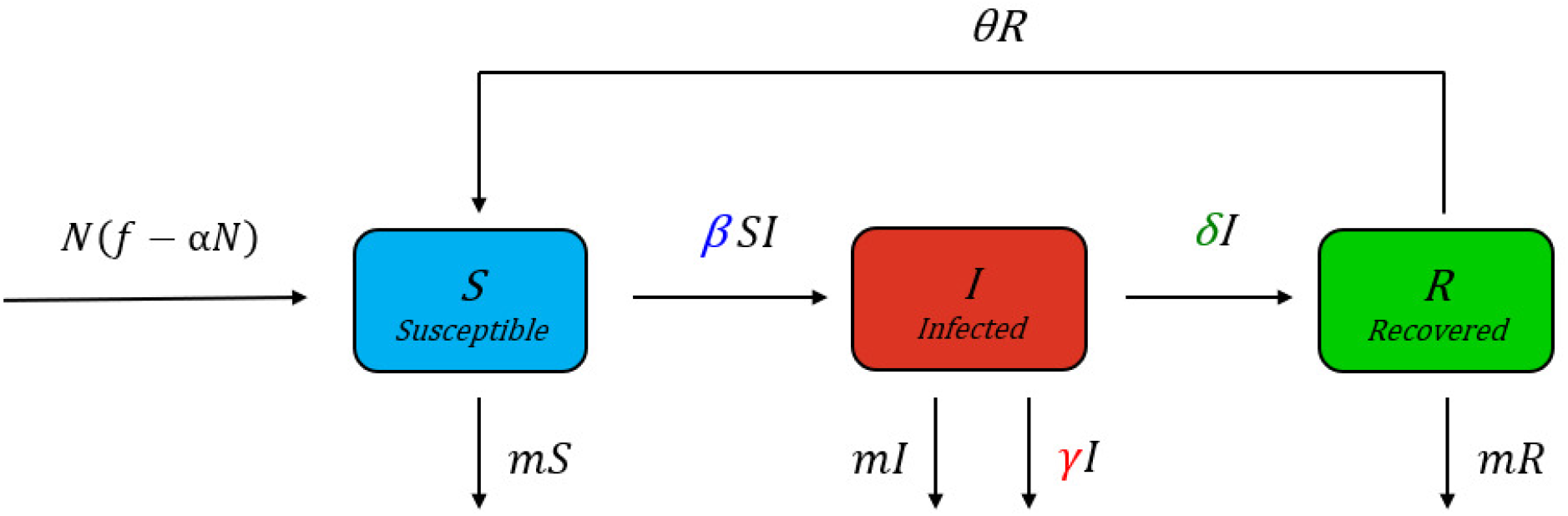
The SIR model. (*S* = susceptible, *I* = infected, and *R* = recovered) given by the set of differential equations (1). The arrows represent the flow of individuals. Parameters colored in blue (host susceptibility *β*), red (pathogen virulence *γ*) and green (host recovery *δ*) are the immunity components that will be considered in the formulation of each fecundity-immunity trade-off.

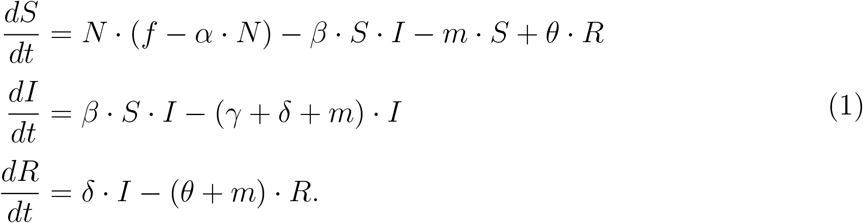

The variable *N* = *S* + *I* + *R* is the total host population, while *S, I*, and *R* denote the susceptible, inflected, and recovered densities, respectively. Usually, in standard SIR models, the input of the susceptible compartment is at a constant rate (Martcheva, 2015). Here, however, we assume that the influx of susceptible individuals follows a logistic growth, as in Gandon et al. (2002) and Best et al. (2009). This logistic growth not only contains an explicit term for host fecundity, which is central in the formulation of the fecundity-immunity trade-off, but also generates a variable population size, which is key in assessing this emergent property in the system. The parameters of this logistic growth include *f*, the *per capita* fecundity of the host, *m*, its natural mortality rate, and *α*, the intraspecific competition within the host population. The pathogen is assumed to be horizontally and directly (contact-based) transmitted, with *β* encapsulating both pathogen infectivity *β*_*y*_ and the *per capita* degree of host susceptibility to infection *β*_*x*_, such that *β* = *β*_*y*_ · *β*_*x*_. However, since only the host is considered to be evolving in this model, *β* only effectively captures variation in host susceptibility, while pathogen infectivity is fixed at unity, such that *β* becomes *β* = *β*_*x*_. The parameter *γ* represents the pathogen virulence, denoting the additional host mortality rate induced by the pathogen. Finally, parameters *δ* and *θ* denote the host recovery rate and the loss of immunity rate, respectively.

This model admits three equilibrium points: a trivial equilibrium (TE), a diseasefree equilibrium (DFE), and an endemic equilibrium (EE). The TE occurs when the abundances of all compartments are zero, which happens only if the fecundity *f* is lower than the natural mortality *m*. We thus always assume that *f > m*, ensuring the longterm maintenance of the host population. In the absence of pathogens (DFE), the host population is fully susceptible and converges to its carrying capacity:

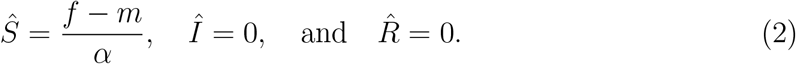

When the pathogen is present in the host population (EE), infected and recovered populations have positive abundances. This equilibrium is given by:

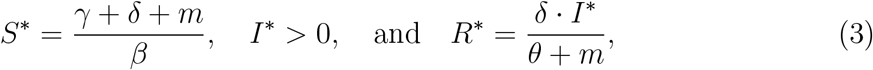

where *I*^∗^, the abundance of the infected individuals, is the solution of a quadratic equation representing the general equation of the endemic equilibrium (see supplementary material S1). The system converges to the DFE when *R*_0_ *<* 1 and to the EE when *R*_0_ *>* 1 (see supplementary figure S1), *R*_0_ being given by:

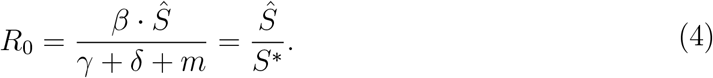

### 2.2 Fecundity-immunity trade-off

We assume that all costs of immunity are expressed as reductions in host fecundity. This is a simplifying assumption —also commonly adopted in other studies—, since immunity costs may, in reality, also affect other life-history traits, such as background mortality or intraspecific competition (Metzler et al., 2023). However, without such simplifications, the analysis of the model would become intractable. In our framework, while fecundity is represented by a single parameter (*f*), immunity encompasses host susceptibility to infection *β*, host recovery *δ*, and host vulnerability to pathogen virulence *γ*. For instance, an immune host can lower its susceptibility to infection, recover faster, or mitigate pathogen virulence (Boots and Bowers, 1999). While we acknowledge that several immune components can be simultaneously activated, here —purely for simplicity—, we will study each of the three immune components separately. We will, however, later discuss the potential implications for host evolutionary outcomes when multiple immune components are jointly activated. These simplifications yield three distinct trade-off relationships: the fecundity-susceptibility, fecundity-recovery, and fecundity-virulence trade-offs. We implement each trade-off by assuming an axis of allocation strategies along which the available resource is constant and uniformly distributed, with optimal positions providing the highest fecundity and the strongest immunity, and where the host’s position *x* determines its phenotype and thus its investment strategy in fecundity versus the corresponding immune component.

In the following, we present the mathematical functions used for the fecundity *f* (*x*), the susceptibility *β*(*x*), the recovery *δ*(*x*), and the virulence *γ*(*x*). Furthermore, although our separation of the three dimensions of immunity is arguably artificial, we propose biological cases that could be close to each of the three immunity scenarios we consider. We then analyze the evolutionary outcome of each trade-off by manipulating its corresponding shape (curvature).

For host fecundity, in all following scenarios, we use a Gaussian-shape curve, given by:

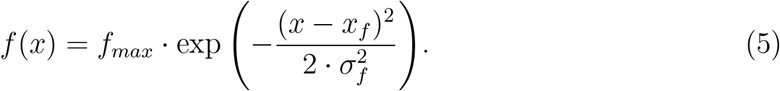

where *f*_*max*_ is the maximum host fecundity in the system, obtained at *x*_*f*_, and *σ*_*f*_ is the width of the fecundity function (figure 2.A). Note that, without any loss of generality, we can always assume that the maximum in fecundity is located at *x*_*f*_, here set at 0 for simplicity, and the width of the fecundity function to be equal to *σ*_*f*_ = 1, simply by rescaling the resource allocation axis.

**Figure 2.**
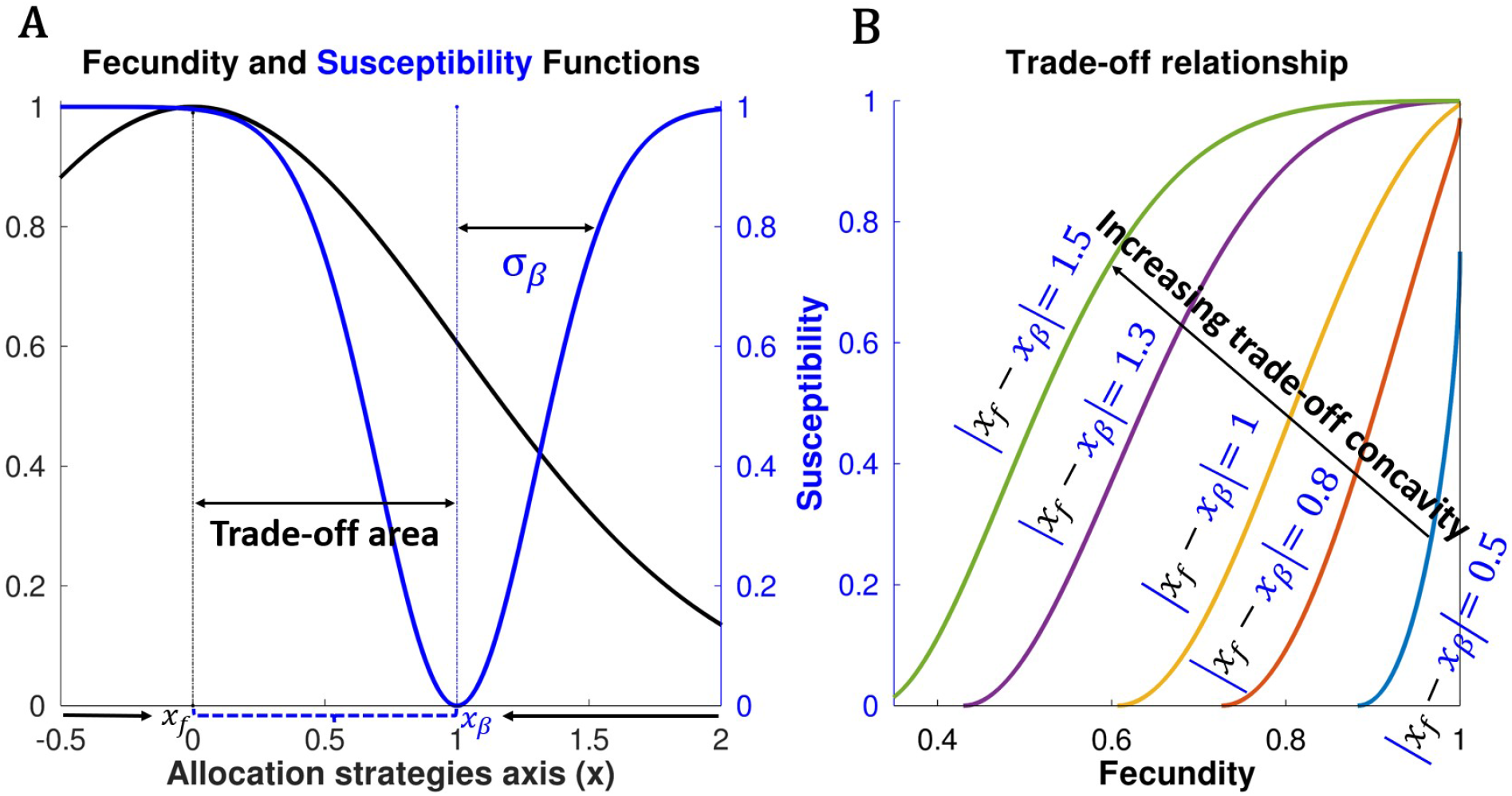
The fecundity-susceptibility trade-off. **Panel A** shows host fecundity (black curve) and susceptibility (blue curve) as functions of host phenotype *x* on the allocation strategies axis. Black and blue vertical lines indicate the position for the optimum in fecundity (*x*_*f*_) and in immunity (*x*_*β*_), respectively. The distance between the optima, |*x*_*f*_ − *x*_*β*_| fixes the trade-off concavity or convexity, as shown on **Panel B.** Parameters: *x*_*f*_ = 0, *σ*_*f*_ = 1, *f*_*max*_ = 1, *σ*_*β*_ = .3, *β*_*max*_ = 1, *q*_*β*_ = 1, *x*_*β*_ = 1 on **Panel A** and *x*_*β*_ = 0.5, 0.8, 1, 1.3 and 1.5 on **Panel B**.

#### 2.2.1 Fecundity-susceptibility trade-off: a zooplankton-fungus system

Our first scenario is inspired by a system in which a zooplankton, *Daphnia dentifera*, becomes infected by a virulent fungus *Metschnikowia bicuspidata*. There are two clones of this zooplankton: the fast-feeding clone and the low-feeding clone. When both clones are uninfected, the fast-feeding clone produces more offspring than the low-feeding clone. However, in the presence of the fungal pathogen, the fast-feeding and high reproductive clone shows a higher risk of fungal infection and produces more fungal spores compared to the low-feeding and less reproductive clone (Hall et al., 2010, 2012), suggesting a fecundity-susceptibility trade-off. To implement this, we build our susceptibility function in such a way that, within the trade-off area, host phenotypes with higher fecundity also exhibit greater susceptibility to pathogen infection compared to phenotypes with lower fecundity. To express the decrease in host susceptibility around the positions maximizing host immunity, we use a downward Gaussian-shaped function given by:

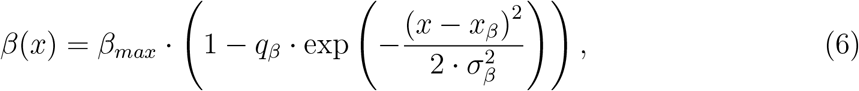

where *x*_*β*_ represents the host phenotype associated with the lowest susceptibility in the system, *β*_*max*_ is the highest possible level of host susceptibility to infection, *q*_*β*_ scales the maximum effect of host immunity in reducing infection, and *σ*_*β*_ is the width of the susceptibility function, with small *σ*_*β*_ leading to relatively higher host susceptibility.

Figure 2 is a graphical illustration representing host fecundity *f* (*x*), susceptibility *β*(*x*), and the trade-off relationship between these two biological functions. While fecundity is optimal at *x*_*f*_, immunity is optimal at *x*_*β*_ where the host susceptibility to pathogen infection is at its lowest level (panel A). Note that only the trait space between these two optima constitutes the effective trade-off area. Evolution necessarily leads to situations within this interval and any singularity will happen therein. Indeed, for *x < x*_*f*_, evolution is directional toward *x*_*f*_, as increases in *x* enhance fecundity while lowering pathogen susceptibility. Likewise, for *x > x*_*β*_, any mutant with a lower *x* has higher fecundity and lower susceptibility to infection, therefore invading. These directional selection dynamics are shown using arrows on figure 2A. In a nutshell, these intervals outside the trade-off area, while biologically debatable, will never be evolutionary accessible. Moreover, the distance set by the separation between the two optima (|*x*_*f*_ − *x*_*β*_|) determines the shape or curvature of the trade-off (panel B).

When the distance is small, the trade-off curve is convex on most of the interval. In this situation, the host can maintain both high fecundity and strong immunity at the same time (blue curve on panel B). In contrast, when the distance is large, the trade-off becomes mostly concave, and achieving high fecundity comes at a substantial cost to immunity (increases host susceptibility) and *vice versa* (green curve on panel B). Note that the investment strategy toward fecundity and immunity is influenced not only by this distance |*x*_*f*_ − *x*_*β*_|, but also by the widths of both the fecundity and susceptibility functions. Here, however, we will assume constant widths of the functions (constant *σ*_*f*_ and *σ*_*β*_), focusing on the distance between the optima. The same rationale applies to the fecundity-virulence trade-off. However, for the fecundity-recovery trade-off, the relationship between the distance |*x*_*f*_ − *x*_*δ*_| and the trade-off curve will be exactly the opposite of what is observed for the two other trade-offs (see supplementary figure S2).

#### 2.2.2 Fecundity-recovery trade-off: a mosquito-protozoan system

Our second scenario could, for instance, be linked to a system in which the female mosquito *Anopheles albimanus* suffers from the malarial parasite *Plasmodium berghei* infection. When the mosquito’s immunity was primed, that is, stimulated, the mosquitoes were able to more easily clear the infection and thus, to recover more easily from the disease compared to unprimed or unstimulated mosquitoes. However, primed mosquitoes that cleared the infection were also less able to produce viable eggs, compared to control or unprimed mosquitoes (Contreras-Garduño et al., 2014), suggesting a fecundity-recovery trade-off. To build our trade-off such that host phenotypes with high fecundity also recover less from the disease, we use the following Gaussian-shaped function for recovery:

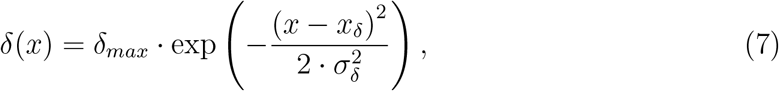

where immunity at position *x*_*δ*_ offers the maximal recovery rate *δ*_*max*_ while *σ*_*δ*_ defines the width of the recovery function (see supplementary figure S2 panels A and B).

#### 2.2.3 Fecundity-virulence trade-off: a beetle-bacteria system

The host immune system can also reduce pathogen virulence —defined as the additional host death induced by pathogens— by limiting pathogen-related damages. A specific example is found in the red flour beetle *Tribolium castaneum* infected by the bacteria *Bacillus thuringiensis*. The immune system of the beetles was primed, and their offspring were later tested by the bacteria. Offspring of primed beetles showed higher post-infection survival, but also endured a reduced early survival, development rate and reproduction (Prakash et al., 2022). These observations suggest the existence of a trans-generational cost of immune priming in terms of reduced fecundity associated with reduced virulence. To model such a fecundity-virulence trade-off, we use a downward Gaussian-shape function for virulence given by:

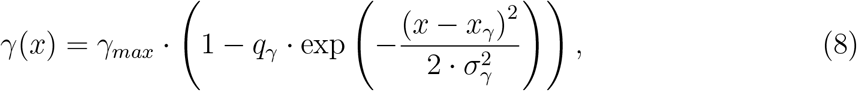

where the host at position *x*_*γ*_ experiences minimal virulence. Parameter *γ*_*max*_ corresponds to the maximum pathogen virulence, *q*_*γ*_ is the maximum reduction in virulence due to host immunity, and *σ*_*γ*_ defines the width of the virulence function (see Supplementary figure S2 panels C and D).

### 2.3 Evolutionary dynamics

We use adaptive dynamics to investigate the consequences of a fecundity-immunity tradeoff on host evolutionary dynamics and on the emergent properties in the evolving system. Adaptive dynamics (Metz et al., 1992; Geritz et al., 1998; Brännström et al., 2013) is a mathematical framework to study eco-evolutionary dynamics, the feedback loop between ecology and evolution, in a given population when one or several quantitative traits are under selection (Dieckmann and Law, 1996; Ferriere and Legendre, 2013). Adaptive dynamics assume the rareness and small effects of the mutations, as well as clonal reproduction and the large size of the focal population (Metz, 2012). Adaptive dynamics therefore differentiate ecological and evolutionary timescales and consider that a population of trait value *x* first reaches its ecological attractor before a rare mutant with trait value *x*_*m*_ appears (Geritz et al., 1997). Under the assumption that the mutant trait *x*_*m*_ is close to the resident trait *x*, the invasion fitness *ω*(*x*_*m*_, *x*), a mathematical function describing the behavior of this rare mutant in its resident population, can be computed. If *ω*(*x*_*m*_, *x*) *>* 0, the mutant trait value increases in frequency until fixation, thus replacing the resident and creating a new environment in which subsequent mutant invasions will be analyzed (Metz et al., 1992; Geritz et al., 1998).

In unstructured populations, invasion fitness is simply defined as the *per capita* growth rate of a rare mutant *x*_*m*_ in the resident *x* population (Metz et al., 1992; Kisdi, 1999). In class-structured populations, however, as in our SIR model, defining invasion fitness is not as straightforward and thus, a Jacobian-based approach has been developed (Gyllenberg and Metz, 2001; Massol et al., 2009; Best et al., 2009; Picot et al., 2019). This Jacobian-based approach takes into account the contribution of each compartment in the mutant invasibility and provides a proxy (a sign equivalent function) of the invasion fitness. This proxy is based on the dominant eigenvalue of the Jacobian matrix of the mutant dynamics when the mutant is rare and the resident is at its ecological equilibrium (full derivation provided in Supplementary Material S2).

It is then possible to follow the direction of host evolution by using the canonical equation of adaptive dynamics, given by:

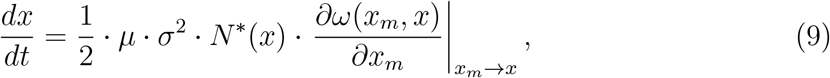

where *µ* is the *per capita* mutation rate, *σ*^2^ is the variance brought by the mutation process, *N* ^∗^(*x*) is the ecological equilibrium of the resident host population expressing trait *x* and *∂ω*(*x*_*m*_, *x*)*/∂x*_*m*_ is the selective gradient (the partial derivative of the invasion fitness (or its proxy) with respect to the mutant trait *x*_*m*_) (Dieckmann and Law, 1996). Interesting phenotypes are those for which the selection gradient vanishes, that is:

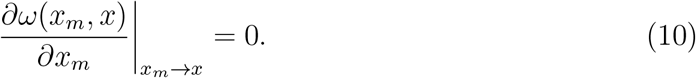

Those points are called singular strategies. Their nature, convergent or not, evolutionary invasible or not, is then determined by the second derivatives of the invasion fitness (Eshel, 1983; Christiansen, 1991; Eshel et al., 1997).

The convergence criterion defines whether phenotypes close to the singular strategy are attracted (convergent) or repelled (non-convergent) by the singularity, whereas the evolutionary stability criterion defines whether the singular strategy can be invaded by a nearby mutant (invasible), or cannot be further invaded (non invasible, ie ESS). Convergent and non-invasible singular strategies are called continuously stable strategies (CSS) and correspond to endpoints of evolution. Convergent and invasible singular strategies correspond to branching points, allowing the evolving host to diverge into two distinct phenotypes (emergence of polymorphism). Finally, non-convergent singular strategies correspond to Repellors or Garden-of-Eden, if they are evolutionary invasible or non invasible, respectively.

## 3 Results

### 3.1 Fecundity-susceptibility trade-off: a zooplankton-fungus system

We find that for a convex trade-offs (low distance between the two stars (optima) on figure 3.A), the host evolves toward a strategy that simultaneously offers maximal fecundity and large immunity, as there is practically no cost associated with investing in one or in the other biological functions. This corresponds to our expectation and such an evolving host eventually clears the disease, leading to maximal host abundances. For a linear tradeoff (, ie a slightly larger distance between the two optima, as shown on figure 3.B), the host displays two alternative and contrasting convergent strategies: (i) a high fecundity strategy (at the expense of immunity), leading to high pathogen propagation (*R*_0_ and the density of infected individuals being larger) and intermediate host abundances; (ii) a second strategy that invests in large immunity at the expense of fecundity, leading to low pathogen propagation, high host abundances, and for which host diversification (the emergence of host polymorphism) is possible. For a concave trade-off (very large distance between the two optima, as depicted on figure 3.C), the cost associated with investing in one or in the other biological function is large. In this case, hosts evolve toward high fecundity at the cost of their immunity. While this strategy leads to an increase in pathogen propagation (large *R*_0_ and increased density of infected individuals), investing in fecundity does not necessarily translate into very large host abundance, which contrasts with our expectations.

**Figure 3.**
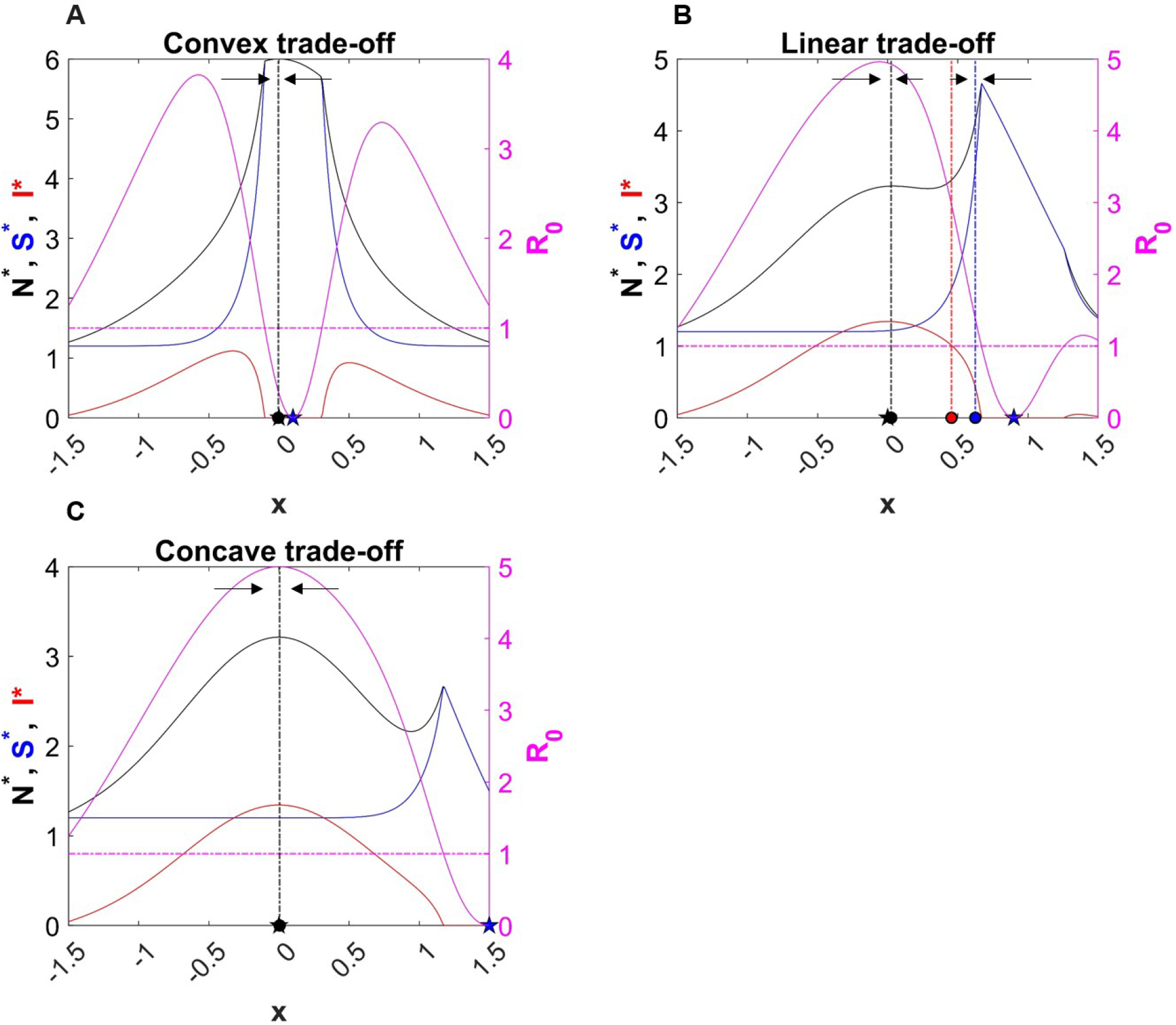
Panel A: Convex trade-off (*x*_*β*_ = 0.10). Solid pink line: disease propagation in the system (*R*_0_), horizontal dashed pink line (*R*_0_ = 1): delimits the regions where the pathogen is present (solid pink line above dashed pink line) from the region where the pathogen is excluded (solid pink line below dashed pink line), solid black line: total host abundance, blue line: density of susceptible individuals, red line: density of infected individuals. The arrows indicate the direction of evolution. On the *x* axis, the black star indicates the trait value for maximum fecundity and the blue star indicates the trait value for minimum susceptibility. The black dot represents the trait value of the singular strategy (CSS), and the vertical dashed black line are used to visualize the position of the CSS relative to disease propagation and host abundance. **Panel B: Linear trade-off (***x*_*β*_ = 0.9). The red dot is the trait value for the Repellor and the blue dot is the trait value for the branching point. The vertical dashed red and blue lines are used to indicate the position of the corresponding singular strategy relative to our two emergent properties. **Panel C: Concave trade-off (***x*_*β*_ = 1.30). Other parameters: *x*_*f*_ = 0, *σ*_*f*_ = 1, *f*_*max*_ = 1, *σ*_*β*_ = 0.3, *β*_*max*_ = 1, *q*_*β*_ = 1, *γ* = 1, *δ* = 0.1, *θ* = 0.1, *α* = 0.15, *M* = 0.1.

While figure 3 is based on one specific set of model parameters, figure 4 provides a broader analysis of how the trade-off shape affects evolutionary outcomes, and highlights that reproduction-oriented strategies are most often selected. Only linear trade-offs lead to an evolution of immunity-oriented strategies. Evolution toward reproduction is observed for convex trade-offs, as immunity and reproduction are then aligned. For concave trade-offs, however, the evolving host invests in fecundity at the expense of immunity, probably because the cost of low fecundity cannot be counterbalanced by high immunity. Linear trade-offs result in moderate investments in immunity, balancing the costs of disease propagation (leading to higher mortality) against the benefits of immediate reproductive output. Note that, on a small interval (from *x*_*β*_ = −1.1 to −0.8, and from *x*_*β*_ = 0.8 to 1.1), the evolutionary outcomes consist in two alternative and convergent singular strategies, a high fecundity and a high immunity strategies. The high immunity strategy is also invasible, i.e., is a branching point that leads to the emergence of host polymorphism (blue region of the curve on figure 3).

**Figure 4.**
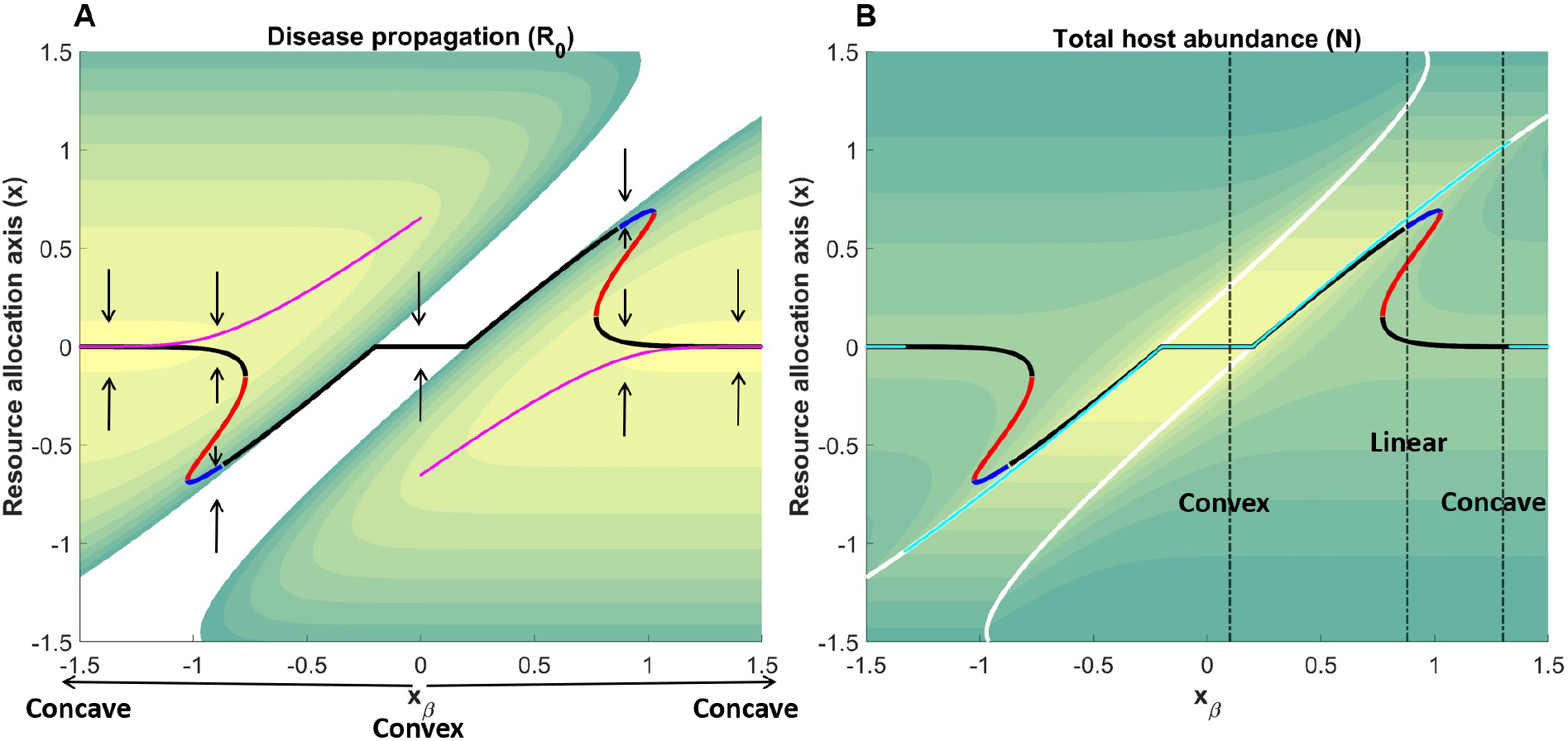
Ecology-Evolution-Environment (E3) diagrams illustrating the impact of the trade-off curvature of a fecundity-susceptibility trade-off, defined by *x*_*β*_, on the host eco-evolutionary dynamics and on disease propagation (panel A) and host abundance (panel B) in the system. **Panel A**: The colored region is the region of host-pathogen coexistence (*R*_0_ *>* 1) while the white region is the region of pathogen exclusion (*R*_0_ *<* 1). Yellow contour colors indicate high pathogen propagation (high *R*_0_) while green contour colors indicate low pathogen propagation (low *R*_0_). Black line: CSS, red line: Repellor, blue line: Branching, Lila line: maximum disease propagation (*R*_0_) in the system for each specific *x*_*β*_. The arrows indicate the direction of evolution. **Panel B**: Yellow contour colors indicate high host abundance (high *N* ^∗^) while green contour colors indicate low host abundance. The cyan line is the maximum host abundance *N* ^∗^ in the system for each specific *x*_*β*_ and the white lines separate the coexistence and the exclusion regions. The dashed black lines show the *x*_*β*_ for the convex, linear and concave trade-offs from figure3. Parameters: *x*_*f*_ = 0, *σ*_*f*_ = 1, *f*_*max*_ = 1, *σ*_*β*_ = 0.3, *β*_*max*_ = 1, *q*_*β*_ = 1, *γ* = 1, *δ* = 0.1, *θ* = 0.1, *α* = 0.15, *M* = 0.1.

These evolutionary dynamics are associated with largely different emergent properties, both in terms of epidemiology and in terms of host abundances. Overall, regarding emergent properties, we observe that, along the singular strategies obtained from a continuous change of the trade-off curve, pathogen propagation *R*_0_ (figure 4.A) and total host abundance *N* ^∗^ (figure 4.B) have opposite directions, that is, for a high pathogen propagation, the total host abundance is low, and *vice versa*. Pathogen exclusion and high host abundance occur under convex trade-offs, while concave trade-offs lead to low host populations and to large epidemics. Indeed, in the second case, evolution favors strategies that are highly vulnerable to pathogen infection, but highly reproductive. Because high reproduction yields more susceptible, this combined with low immunity favors pathogen propagation (higher *R*_0_). Interestingly, in the region ranging from *x*_*β*_ = −0.8 to −0.5, and from *x*_*β*_ = 0.5 to 0.8, the selected strategies are located at the edge of the hostpathogen coexistence domain (*R*_0_ very close to 1). However, being in these regions still allows the host to have a sufficiently good fecundity. These regions therefore constitute, for the host, a good compromise between reducing pathogen harm (pathogen infection) and having a good fecundity. Because pathogens are already close to extinction under these conditions in our deterministic framework, we anticipate that including stochastic components (e.g., demographic stochasticity) in such conditions may further promote parasite exclusion, thus expanding the domain in which the pathogen is excluded.

### 3.2 Fecundity-recovery trade-off: a mosquito-parasite system

The host evolutionary outcome in a fecundity-recovery trade-off is qualitatively similar to the evolutionary outcome that we observed with a fecundity-susceptibility trade-off (see Supplementary figure S3). The explanation behind this similarity could be that reducing host susceptibility to infection or increasing host recovery both result in qualitatively similar states from a population point of view; a higher investment in immunity leads to a decreased abundance of infected individuals and to an increase of susceptible individuals. Since infected individuals determine disease incidence, reducing their density would, in turn, decrease pathogen propagation in the population. We can also mathematically explain this similarly by studying the equation of the basic reproduction number *R*_0_ (equ. 4). One can notice that decreasing *β* or increasing *δ* would have qualitatively similar effects on *R*_0_, decreasing its value. Both biological and mathematical explanations are comparable and may, in the fecundity-susceptibility and fecundity-recovery trade-offs, result in structuring the host population similarly.

### 3.3 Fecundity-virulence trade-off: a beetle-bacteria system

The evolutionary outcome for a fecundity-virulence trade-off is partially qualitatively different from the two other trade-offs investigated. Figure 5 shows that under a convex fecundity-virulence trade-off, the host evolves toward a strategy investing at the same time in fecundity and in immunity (reducing pathogen virulence). Evolution then maximizes host abundance. Such evolutionary strategies also lead to higher pathogen propagation, contrary to what is observed with the fecundity-susceptibility and fecundity-recovery trade-offs, which is inconsistent with our expectations. This outcome can however be explained, on the basis on basic biological components. Here, selection for immune strategies extends the lifespan of infected individuals, leading to an increase in their density, and to higher disease propagation within the population. Equation for *R*_0_ confirms this (equ. 4), as decreasing *γ* would inevitably result in increased disease propagation. The same reasoning applies to concave trade-offs where evolution favors fecundity at the expense of immunity, leading to a decrease in disease propagation. Indeed, under concave trade-offs, selection for fecundity inevitably increases pathogen-induced death rate, shortening the lifespan of infected individuals and reducing their density as well as disease propagation. These concave trade-offs also result in an increase in host abundance, probably because selection for high fecundity is, in this case, able to counterbalance the negative effect of having bad immunity. For linear trade-offs, the evolutionary dynamics exhibit two alternative outcomes, either (i) a strategy investing in fecundity at the expense of immunity, with relatively lower pathogen propagation and higher host abundance, or (ii) a second strategy investing in immunity at the expense of fecundity, with higher pathogen propagation and lower host abundance. For the investigated set of parameters and regardless of the trade-off curve, host diversification is not possible, and the evolving host is unable to get rid of the pathogen throughout evolution.

**Figure 5.**
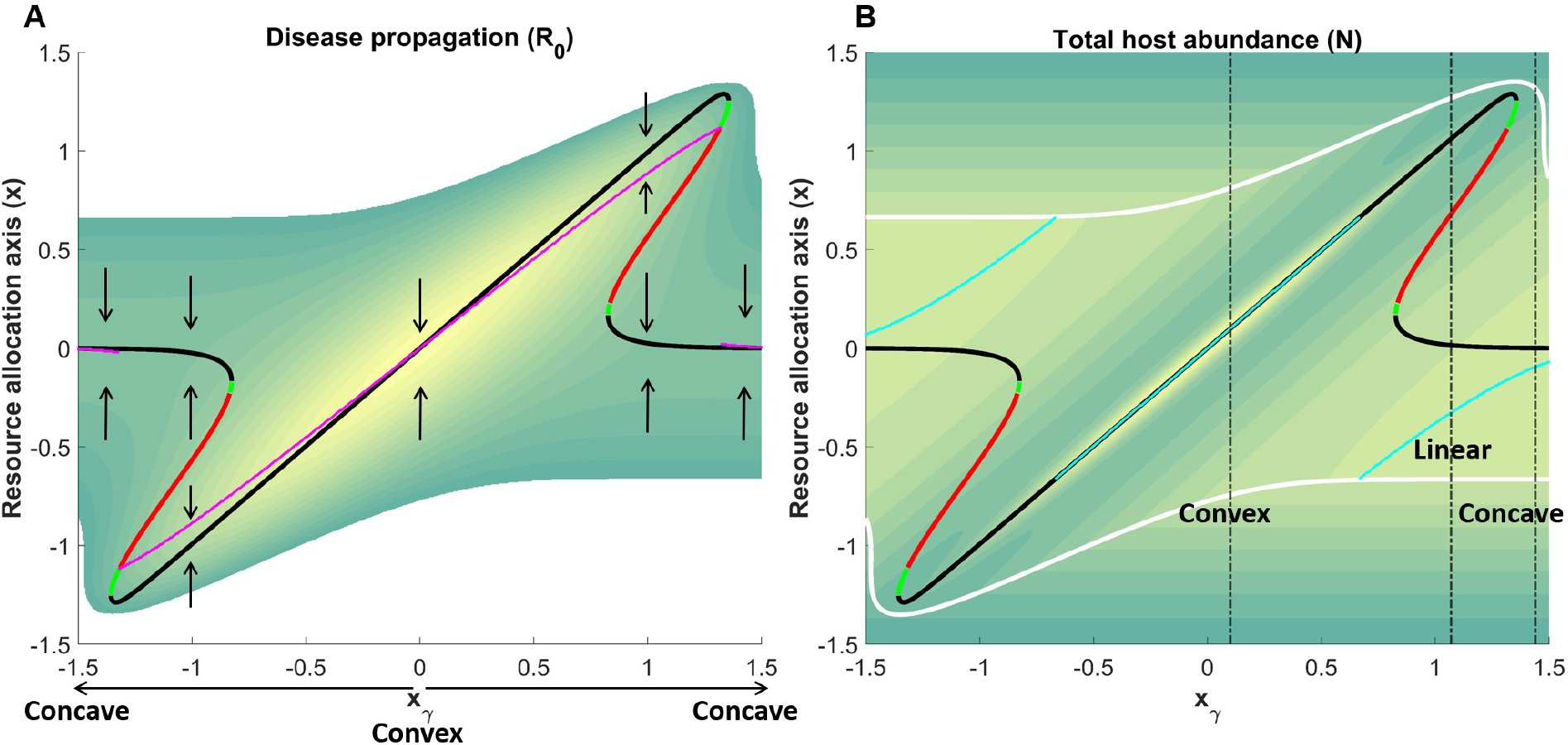
Ecology-Evolution-Environment (E3) diagrams illustrating the impact of the trade-off curvature (*x*_*γ*_) on the host eco-evolutionary dynamics and on disease propagation (panel A) and host abundance (panel B) in the system. Yellow contour colors indicate high pathogen propagation and high host abundance on the respective panels, while green contour colors indicate low pathogen propagation and low host abundance. **Panel A**: The colored region is the region of host-pathogen coexistence (*R*_0_ *>* 1) while the white region is the region of pathogen exclusion (*R*_0_ *<* 1). Black line: CSS, red line: Repellor, green line: Garden of Eden, Lila line: maximum disease propagation (*R*_0_) in the system for each specific *x*_*γ*_. The arrows indicate the direction of evolution. **Panel B**: The cyan line is the maximum host abundance (*N* ^∗^) in the system for each specific *x*_*γ*_ and the white lines delineate the boundary between the coexistence and the exclusion regions. The dashed black lines show the *x*_*γ*_ for a convex, linear and concave trade-offs. Parameters: *x*_*f*_ = 0, *σ*_*f*_ = 1, *f*_*max*_ = 1, *σ*_*γ*_ = 0.4, *γ*_*max*_ = 40, *q*_*γ*_ = 1, *β* = 10, *δ* = 20, *θ* = 30, *α* = 0.1, *M* = 0.2.

**Figure 6.**
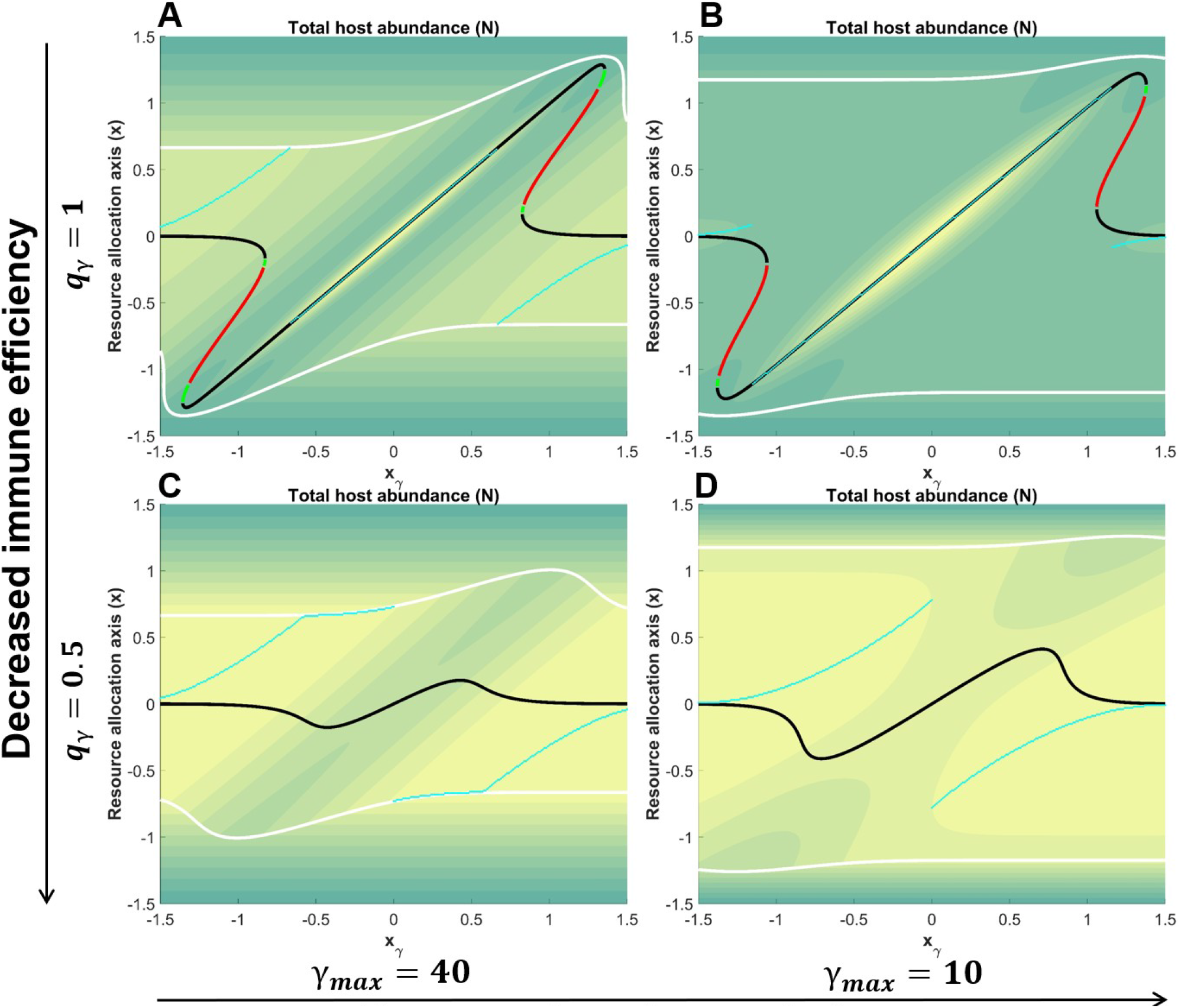
This figure shows the impact of different immune efficiency *q*_*γ*_ and maximum virulence *γ*_*max*_ on host abundance in the system. Yellow contour colors indicate high host abundance on the respective panels, while green contour colors indicate low host abundance. Parameters: *x*_*f*_ = 0, *σ*_*f*_ = 1, *f*_*max*_ = 1, *σ*_*γ*_ = 0.4, *β* = 10, *δ* = 20, *θ* = 30, *α* = 0.1, *M* = 0.2.

While these results assume a perfectly efficient immune response and a relatively high maximum virulence, they still hold for disease propagation when such constraints are relaxed, but not for the variations in host density. Indeed, when the immune efficiency *q*_*γ*_ decreases, high host abundances are observed primarily under concave trade-offs. Conversely, when the maximum virulence *γ*_*max*_ decreases, high host abundances are observed mainly under convex trade-offs. This suggests that the interplay between the efficiency of the host immune system and the intensity of pathogen virulence, which do not affect the trade-off shape, may however influence host abundance, at least for this specific immune component investigated.

## 4 Discussion

In an attempt to disentangle the eco-evolutionary outcome in a host-pathogen system under resource allocation constraints between fecundity and immunity, we found that when there is no or little cost associated with investing in fecundity or immunity, then the host evolutionary strategy is to invest in both functions. As the cost increases to an intermediate level such that investing in one biological function has a moderate cost in the other function, the host displays two alternative and convergent singular strategies (one toward immunity and the other toward fecundity). Furthermore, for the fecundity-susceptibility and fecundity-recovery trade-offs, the alternative and convergent singular strategy investing into immunity sometimes leads to a branching point that allows the emergence of polymorphism. Finally, when the trade-off cost becomes very high, investing in one biological function is highly costly for the other function. In this case, the host evolves increased fecundity despite the damages it may suffer from reduced immunity. We also observed along the singular strategies, the points toward which host evolution is attracted or from which evolution is repelled, that in general, for the fecundity-susceptibility and fecundity-recovery trade-offs, pathogen propagation (*R*_0_) negatively affects host abundance such that when pathogen propagation is high, host abundance is low, and *vice versa*. However, for the fecundity-virulence trade-off, an increase in *R*_0_ does not necessarily correlate with a decrease in host abundance along evolutionary trajectories, as host abundance in this case seems to be more sensitive to other model parameters such as maximum pathogen virulence and host immune system efficiency.

Although in an immune host reducing susceptibility to infection *β* or increasing recovery *δ* are two distinct defense mechanisms, they entail similar host evolutionary outcomes. We explain this similarity by the fact that both mechanisms are resistance mechanisms able to reduce the density of infected individuals, resulting in higher frequencies of susceptibles. As infected individuals undergo two different death rates, the natural one and the pathogen-induced one (virulence), any mechanism that selects for strategies with low density of infected individuals will also, by extension, favor a reduction in the total number of pathogen-induced deaths. Consequently, the loss of host abundance in the system will slow down, leading to the maintenance of a relatively higher host density. This may explain the observed antagonistic relationships between disease propagation and total host abundance along evolutionary trajectories, as an indirect consequence of changes in the density of infected individuals —either increasing or decreasing— as the host evolves toward higher fecundity or greater immunity, respectively. We stress that this negative link between disease propagation and host density is distinct from the well-expected direct epidemiological effect. The relationship we report here is to be understood along evolutionary trajectories. I.e., in addition to the directly negative effects of pathogens, natural selection can select phenotypes that further decrease populations size. However, hosts expressing sufficiently strong immune responses can counter these negative pathogen effects and maintain relatively high population densities. In the extreme cases, these immune hosts are even able to reduce disease propagation and thus, the density of infected individuals, under the critical level of disease maintenance in the population (*R*_0_ *<* 1). The resulting consequence is the pathogen exclusion from the system, leading the host to have the highest abundance (see figure 4 and figure S3.A and B).

Contrary to this, when the defense mechanism implies a reduction of pathogen virulence *γ*, infected individuals experience an increase in their survival, resulting in an increase in disease propagation along evolutionary trajectories. However, regarding the other emergent property, the dynamics is less straightforward as what is observed with the two first trade-offs, as immune efficiency and maximum virulence seem to also influence the total host abundance, irrespective of the trade-off shape. Concretely, host evolution with a less efficient immune system leads to an increase in host abundance under concave trade-offs, while evolution with a less virulence pathogen results in an increase of host abundance under convex trade-offs. The difference observed between the fecundity–virulence trade-off and the two other defense strategies stems from the specific nature of virulence reduction, which operates through a distinct immune mechanism. Indeed, this defense mechanism can be viewed as a tolerance mechanism, since the host strategy is to carry the infection and to reduce pathogen-associated deaths. Resistance and tolerance mechanisms have already been shown to lead to different evolutionary outcomes in different host-pathogen systems (Svensson and Råberg, 2010). Vitale and Best (2019) showed that, when resistance carries no cost, its evolution drives the trait toward maximization, ultimately causing parasite extinction. In contrast, tolerance does not lead to parasite elimination because of the positive feedback it generates with infection prevalence, perfectly matching our own observations. Furthermore, field studies of diverse plant species infected by rust fungi confirm that resistance traits tend to be polymorphic, and tolerance traits tend to be fixed (Roy and Kirchner, 2000), while in other host-pathogen systems, tolerance may prevent host castration and resistance may result in the complete loss of fecundity in infected hosts (Best et al., 2010). Therefore, it is not surprising that this difference between resistance and tolerance mechanisms observed in these previous studies, in the way they affect host evolutionary dynamics, may account for the way they differently affect emergent properties in the system, as detected here.

Although our results demonstrate the impact of resource allocation constraints in host-pathogen interactions over the course of evolution, they should be interpreted with caution. Indeed, our simplified evolutionary scenarios —where fecundity is linked to each immunity component separately— may lead to misleading conclusions if, in reality, multiple immune components interact. While this simplification is justified by the fact that trade-offs involving fecundity and two or more immunity components may render our analysis considerably more complex, fecundity-immunity trade-offs can be far more intricate than this simple version presented here. In particular, the activation of the immune response may require the simultaneous engagement of multiple immune mechanisms (Bonneaud et al., 2019), with the cost associated with each of them. For example, in humans, the multidimensional nature of immune trade-offs can be illustrated by the balance between innate and acquired immunity, such that investment in one component may constrain investment in the other, and this balance is thought to be optimized in response to local ecological conditions, including nutritional abundance at the early developmental stage (McDade et al., 2016; Georgiev et al., 2016). The same allocation trade-off between innate versus acquired immunity has also been reported in birds (Minias et al., 2023), in reptiles (Sandmeier et al., 2012), and in fishes (Wegner et al., 2007), suggesting that both mechanisms can be jointly activated, but at different levels. It is therefore possible for an infected host to fragment and allocate the available resource to different immunity mechanisms or components at the same time, if this strategy offers a better selective advantage over investing the total available resource in only one immunity mechanism or component (Bonneaud et al., 2019). This questions the evolutionary outcome in host-pathogen systems, where more complicated trade-offs with more than one immunity trait are incorporated. In our case, this would imply an infected host partitioning and allocating the available resource to fecundity and to all immunity components or mechanisms (reduced susceptibility, enhanced recovery, and reduced virulence) at the same time. Assuming there are sufficient energetic resources that are equally allocated among immune mechanisms, we hypothesize that the combined effects of reduced susceptibility and increased recovery (i.e., resistance mechanisms) may exceed the effect of reduced virulence (i.e., a tolerance mechanism), as resistance and tolerance exhibit different evolutionary outcomes. In this case, we could expect the evolutionary outcome to be similar to the one we observed in a fecundity-susceptibility or in a fecundity-recovery trade-off.

While it is already admitted that host-pathogen systems may exhibit complex dynamics, our results suggest that, a particular resource allocation constraint, a fecundity-immunity trade-off, may alter some emergent properties in the system as well as the host evolutionary outcome. These findings could constitute a contribution to the understanding of the long run disease spread and host productivity (abundance) in such systems, as infected host are usually engaged in costly defense mechanisms against invading pathogens. Interestingly, under specific conditions, host diversification is possible, raising questions not only about the long run maintenance of such diversity, but also about the maximum coexisting phenotypes the system can support, thereby motivating further investigations. While this current study has been carried out by considering host evolution alone, we believe that a co-evolutionary approach is needed, in order to apprehend how pathogens could evolutionary respond to host evolution and how, in this case, the evolutionary dynamics in the system may differ from the evolutionary dynamics when host alone is evolving.

## Acknowledgments

This work was founded by the Swiss National Science Foundation Project grant 31003A-182386. RPR also acknowledges funding from the Swiss National Science Foundation Sinergia grant CRSII5-202290, and a lead-agency grant 320030L-227556.

## Author contributions: CRediT

**Edgard B. Djahoui**: Conceptualization, Methodology, Formal analysis, Writing - Original Draft. **Nicolas Loeuille**: Conceptualization, Writing - Review & Editing, Supervision. **Rudolf P. Rohr**: Conceptualization, Writing - Review & Editing, Supervision, Funding acquisition.

## Supplementary material

### S1 Ecological analysis

The ecological dynamic is given by:

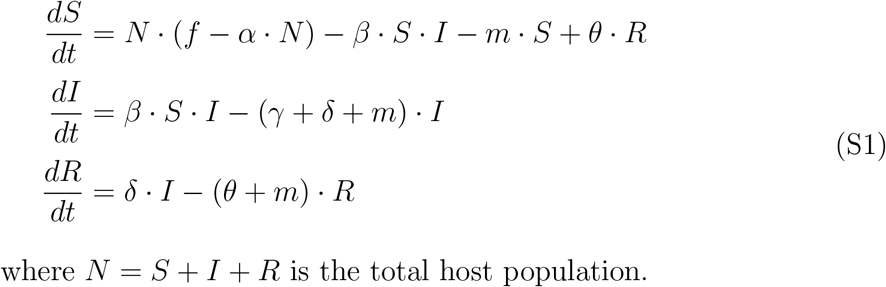

The model admits 3 equilibrium points. The first two ones, the trivial and the disease free ones, have already been presented in the main text. Here we will present details concerning the third equilibrium, the endemic one. To find the feasible solution of this equilibrium, we need to solve the following system:

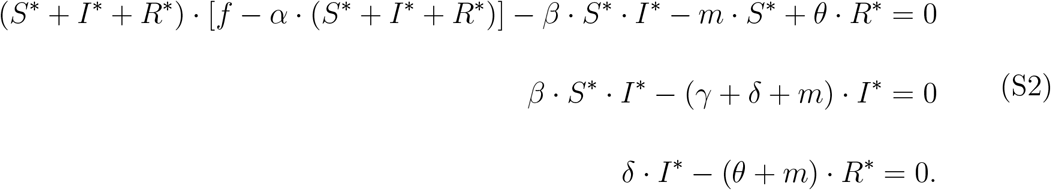

From the second and third equations of S2, it is easy to extract *S*^∗^ and *R*^∗^ as:

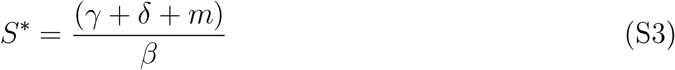

And

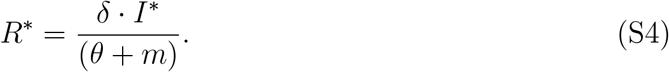

Replacing S4 in the first equation of S2 gives:

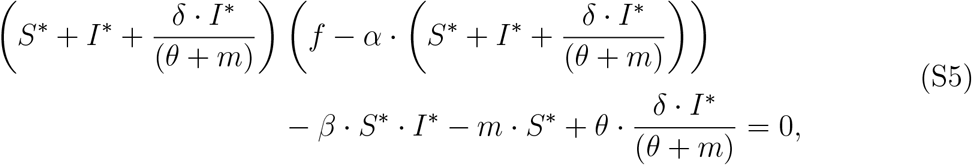

which after some minor algebra transformations, becomes:

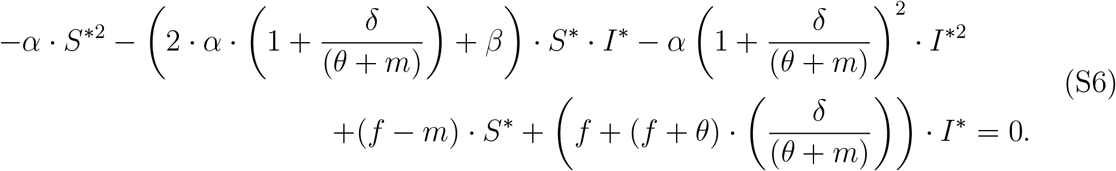

This is the general equation of the endemic equilibrium, in *S*^∗^ and *I*^∗^. Equation S6 is the equation of a conic *C* for which the discriminant Δ is:

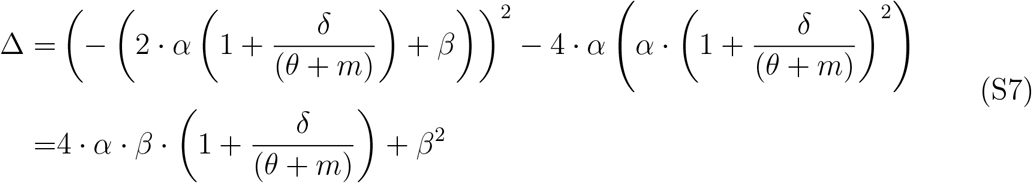

As all parameters are positive, the discriminant of the conic *C* is positive (Δ *>* 0) and, thus, it is always a hyperbola. Moreover, we have

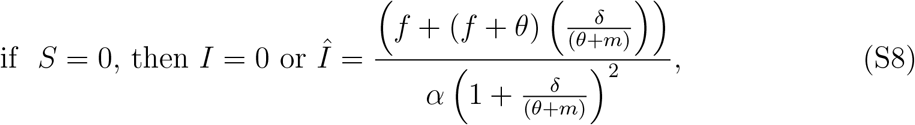

And

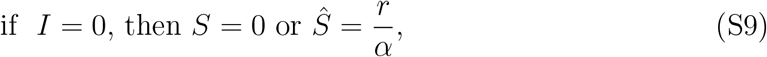

which imply that only one branch of the conic *C*, the superior branch, crosses the border of the positive domain, the domain at which *S >* 0 and *I >* 0, at (*Ŝ*, 0) and at (0, *Î*) (blue curve on figure 4). Therefore, there is only one positive equilibrium point, localized at the point where *S*^∗^ crosses the superior branch of the conic *C* (red dot of figure 4). Moreover, *S*^∗^ crosses the superior branch of the conic *C* if and only if *S*^∗^ *< Ŝ*, which is equivalent to *R*_0_ *>* 1:

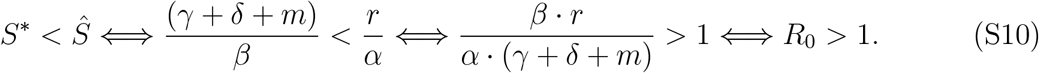

We then conclude that there exists one endemic equilibrium only if *R*_0_ *>* 1, and given by

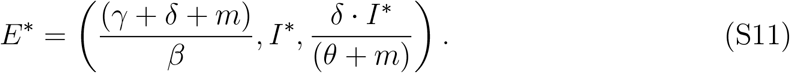

with *I*^∗^ *>* 0 the positive solution of the quadratic equation S6

In our work, *I*^∗^ was numerically calculated by replacing *S*^∗^ (equation S3) into equation S6, and then solving the quadratic equation S6, thus, by finding the positive solution of *C* under the condition *R*_0_ *>* 1.

### S2 Jacobian-based invasion criterion

The dynamic of the mutant host *x*_*m*_ when rare in a resident population *x* is given by:

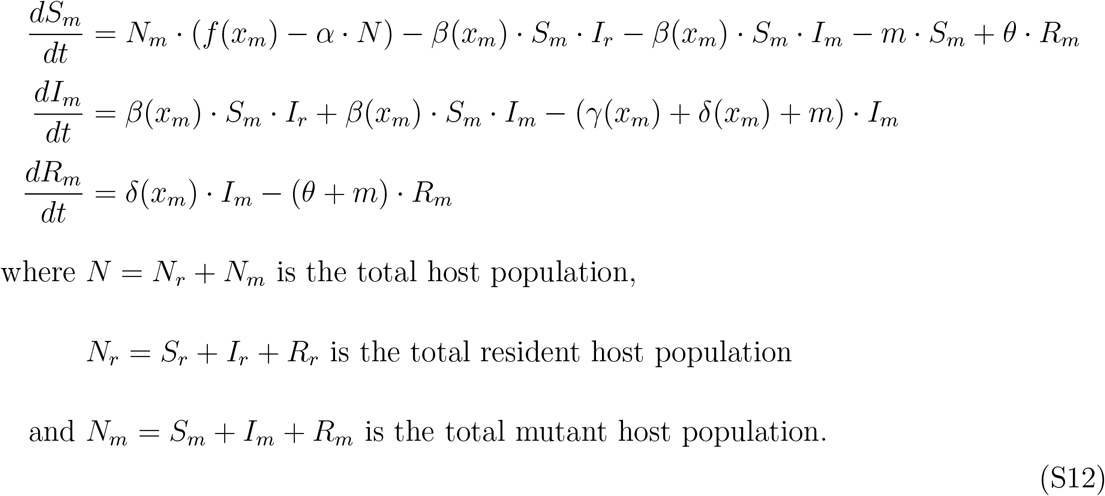

The Jacobian matrix *J* of this mutant host dynamics around the resident host assumed to be at equilibrium is given by:

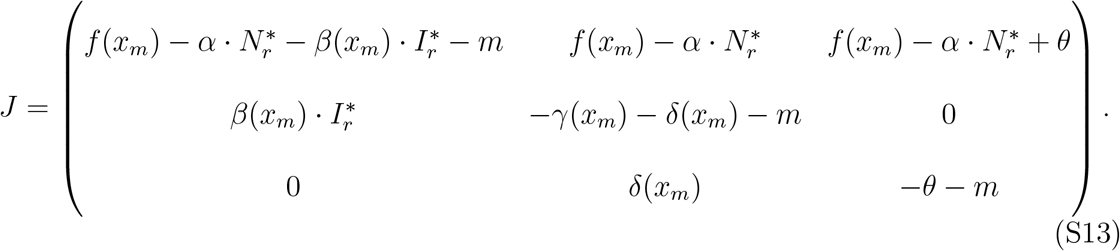

We recall that this assumes the mutant is so rare at the beginning of the invasion that its density is negligible. We analyzed the stability of *J* by finding its eigenvalues *λ, id es* by solving *P* = det(*J* − *λI*) = 0, where *I* is the identity matrix. The real part of the dominant eigenvalue *λ*_*d*_ (the largest one) of *P* provides the proxy for the invasion fitness. If the real part of *λ*_*d*_ is positive, then *J* is unstable, and the mutant host can invade and replace the resident host in the evolutionary course.

## S0.1 Supplementary figures

**Figure S1.**
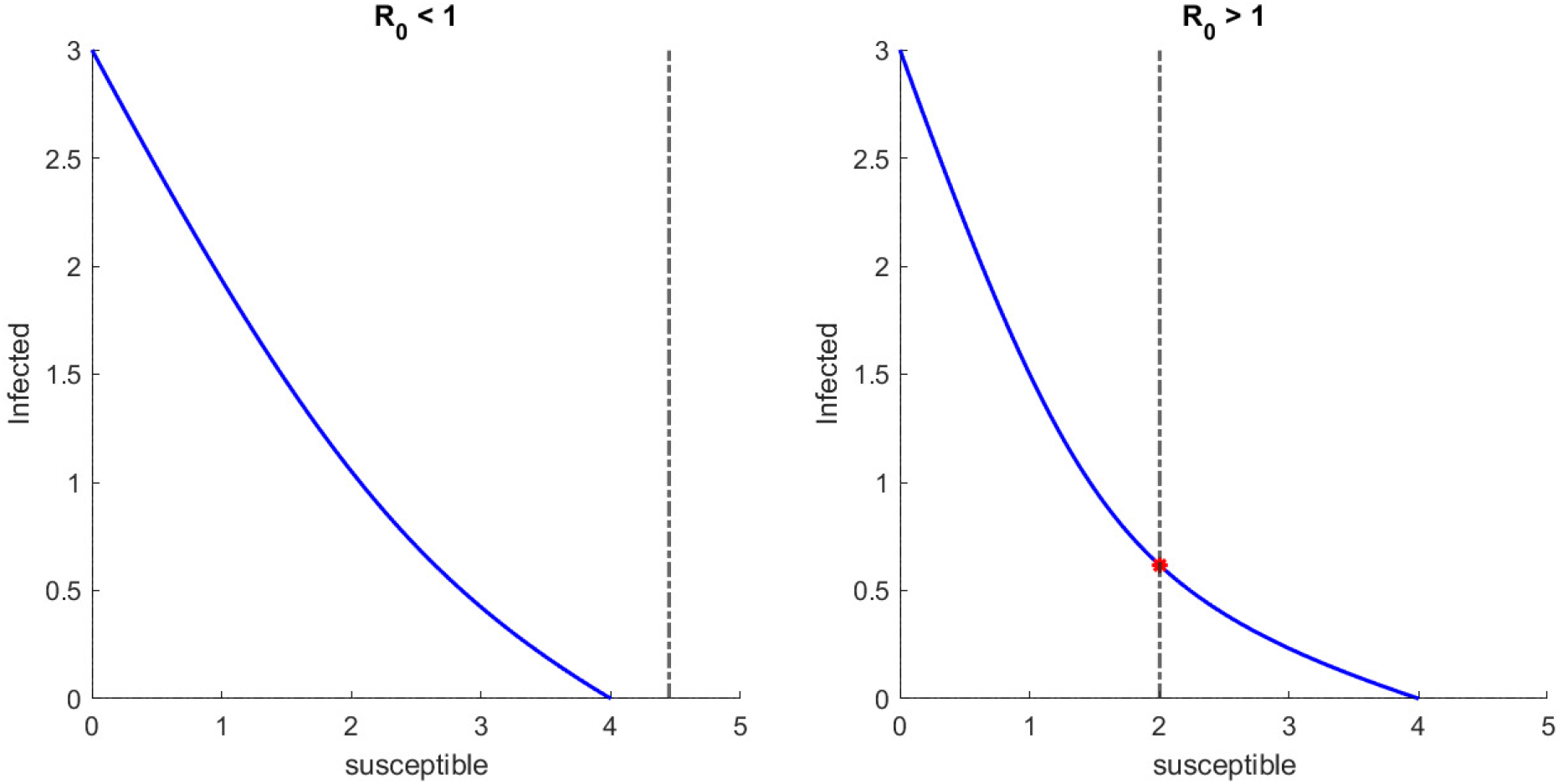
**Panel A** shows the case of a disease free equilibrium, while **panel B** illustrates an endemic equilibrium. On both panels, the blue curve is the superior branch of the conic *C* and the black dashed vertical line represents the density of the susceptible individuals at the endemic equilibrium. An endemic equilibrium exists with positive solution of both susceptible and infected individuals if and only if the blue curve crosses the black dashed vertical line in the positive region of the domain (red point of panel B). Parameters: *f* = 2, *M* = 0, *α* = 0.5, *γ* = 2, *δ* = 2, *θ* = 2, *β* = 0.9 for panel A and *β* = 2 for panel B.

**Figure S2.**
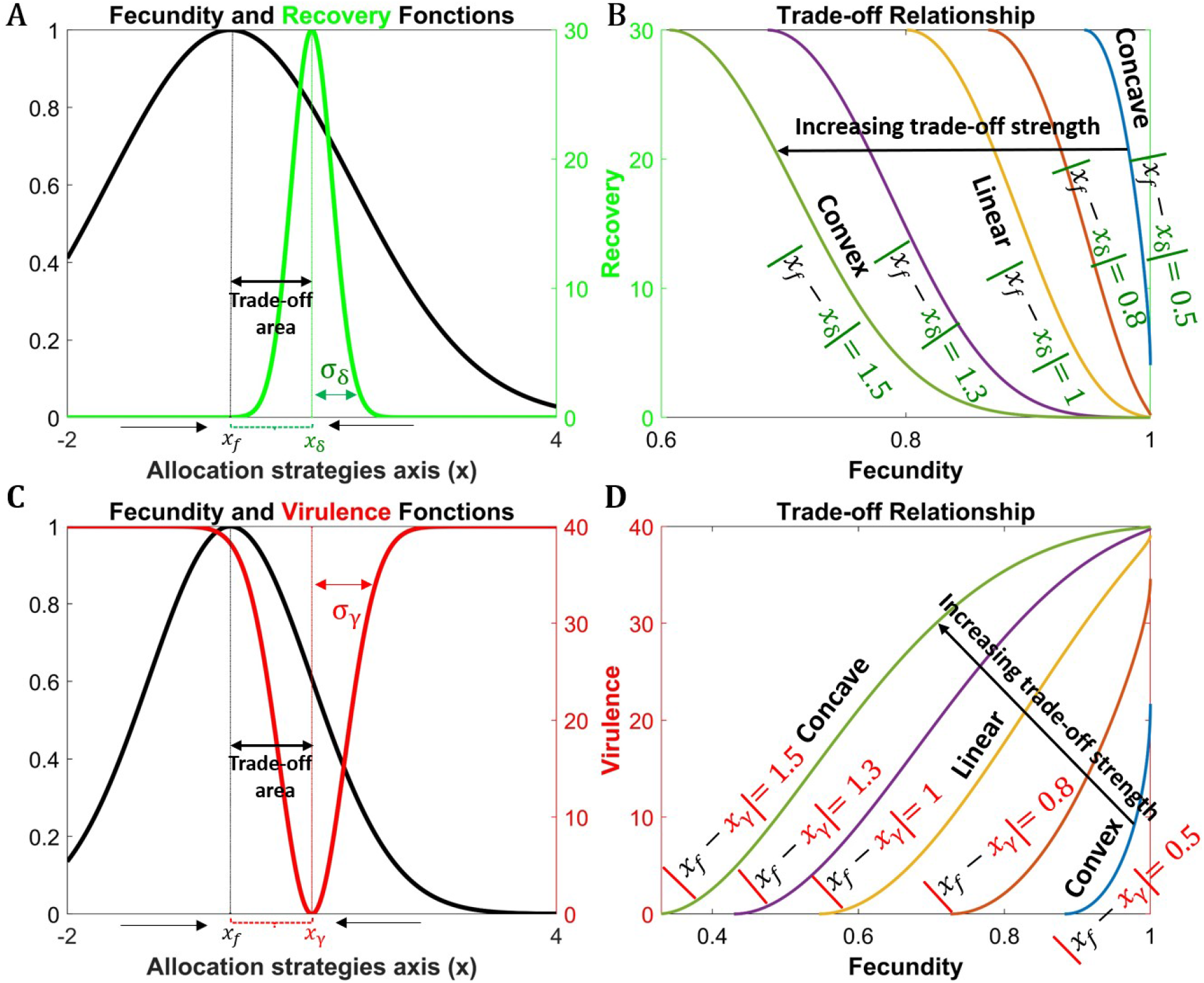
Panel A and panel B illustrate the fecundity-recovery trade-off, while Panel C and panel D illustrate the fecundity-virulence trade-off. **Panel A** represents the fecundity (black curve) and recovery (green curve) as function of the host position on the resource allocation axis *x*. The black and green vertical lines indicate the optimum in fecundity (*x*_*f*_) and in recovery (*x*_*δ*_), respectively. The distance between the optima |*x*_*f*_ − *x*_*δ*_ | determines the trade-off curve and strength. **Panel B** represents the relationship between the fecundity and recovery functions for various distances between the optima of the two functions, that is, for various trade-off strengths. Parameters: *x*_*f*_ = 0, *σ*_*f*_ = 1.5, *f*_*max*_ = 1, *x*_*δ*_ = 1, *σ*_*δ*_ = 0.25, *δ*_*max*_ = 30. **Panel C** shows the fecundity and virulence (red curve) as function of the host position on the resource allocation axis *x*. The red vertical line indicates the optimum in immunity (*x*_*γ*_), where pathogen virulence is reduced to its lowest value in the system. The distance between the optima |*x*_*f*_ − *x*_*γ*_ | determines the trade-off curve. **Panel D** illustrates the relationship between the fecundity and virulence functions for various fecundity-virulence trade-off strengths. Parameters: *x*_*f*_ = 0, *σ*_*f*_ = 1, *f*_*max*_ = 1, *x*_*γ*_ = 1, *σ*_*γ*_ = 0.4, *γ*_*max*_ = 40, *q*_*γ*_ = 1.

**Figure S3.**
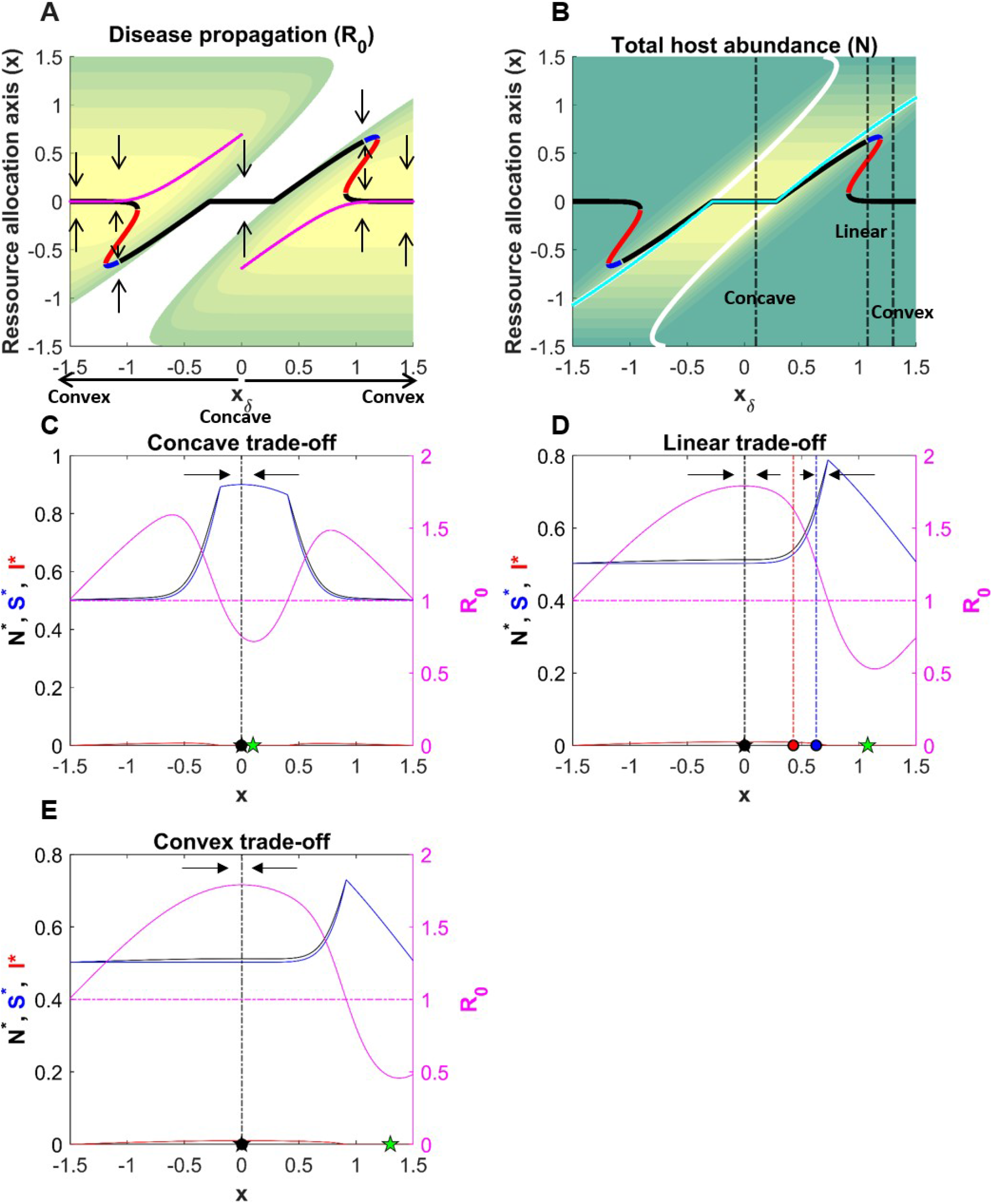
Panel A and panel B are E3 diagrams illustrating the impact of the trade-off curve of a fecundity-recovery trade-off, set by *x*_*δ*_, on the host eco-evolutionary dynamics and on disease propagation (panel A) and host abundance (panel B) in the system. **Panel A**: The colored region is the region of host-pathogen coexistence (*R*_0_ *>* 1) while the white region is the region of pathogen exclusion (*R*_0_ *<* 1). Black line: CSS, red line: Repellor, blue line: Branching, Lila line: maximum disease propagation (*R*_0_) in the system for a given *x*_*δ*_. The arrows indicate the direction of evolution. **Panel B**: The cyan line is the maximum host abundance (*N* ^∗^) in the system for a given *x*_*δ*_ and the white line indicates the transition between the coexistence and the exclusion regions. The dashed black lines show the *x*_*δ*_ for the concave, linear and convex trade-offs that are represented in the 3 panels below. Parameters: *x*_*f*_ = 0, *σ*_*f*_ = 1.5, *f*_*max*_ = 1, *σ*_*δ*_ = 0.25, *δ*_*max*_ = 30, *β* = 40, *γ* = 20, *θ* = 2, *α* = 1, *M* = 0.1. **Panel C: Concave trade-off (***x*_*β*_ = 0.10). Solid pink line: disease propagation in the system (*R*_0_), horizontal dashed pink line (*R*_0_ = 1): delimits the regions where the pathogen is present (solid pink line above the dashed pink line) from the region where the pathogen is excluded (solid pink line below the dashed pink line), solid black line: total host abundance, blue line: density of susceptible individuals, red line: density of infected individuals. The arrows indicate the direction of evolution. On the *x* axis, the black star indicates the trait value for maximum fecundity and the green star indicates the trait value for maximum recovery. The black dot is the trait value of the singular strategy (CSS) and the vertical dashed black line helps visualize the position of the CSS relative to the emergent properties. **Panel D: Linear trade-off (***x*_*δ*_ = 1.077). The red dot is the trait value for the Repellor and the blue dot is the trait value for the branching point. The vertical dashed red and blue lines help visualize the position of the corresponding singular strategy relative to the emergent properties. **Panel E: Convex trade-off (***x*_*δ*_ = 1.30).

